## Supplementary note and figures for "Linked-read sequencing of gametes allows efficient genome-wide analysis of meiotic recombination"

Sun and Rowan *et al*

#### Supplementary Note 1: CO identification within simulated data

To adjust our CO identification pipeline, we performed a simulation analysis. For this we used the reference sequences of *Arabidopsis thaliana* Col-0^1^ and L*er*^2^ to simulate a pool of non-recombinant genomes from both accessions (similar to sequencing non-recombinant F1 hybrids). Sixty million read pairs were simulated for each of the parental genomes using *LRSim*^3^ by setting the number of barcodes to ~1.35 million (each with 3 molecules) and the maximum molecule size to 65 kb. There were 4,399,970 molecules recovered with short read alignments to the Col-0 genome and 4,382,340 molecules with short read alignments to the L*er* genome. Molecule characteristics recovered after short read alignments were generally comparable to the simulated molecules before read simulations (Supplementary Figure 8a-b).

Though most of the recovered molecules were shorter as the maximal simulated molecule size, there were some recovered molecules, which were longer than 65 kb. These resulted from the co-occurrence of independent molecules from adjacent genomic regions with the same barcode. Such chimeric molecules (Supplementary Figure 8c) can lead to a (false) recombinant molecule prediction if the two falsely merged molecules come from different parental genomes. We therefore included a maximum molecule length cut-off for the recovered molecules, which can be adjusted according to specific library characteristics.

However, even after filtering the simulated data for the maximum molecule length of 65 kb, we still detected 34 recombinant molecules using the Col-0 genome as reference sequence and 10 recombinant molecules with the L*er* genome. All of these false CO predictions resulted from ambiguous alignments of L*er* reads to the Col-0 genome or Col-0 reads to the L*er* genome. All false recombinant molecules in Col-0 reference-based analysis were different from those in L*er* reference-based analysis, and thus all such FPs could be filtered by intersecting the two sets. As a consequence, we also applied this filtering for analyzing the real sequencing data and only kept those recombinant molecules that were present in analysis using both parental reference sequences.

#### Supplementary Note 2: Overlapping recombinant molecules

*Overlapping recombinant molecules from identical CO events*

As the number of distinct, haploid genomes *G* in a pool is typically limited, there can be more than one recombinant molecule covering the same CO (as the different recombinant molecules can be sampled from the same site from two different genomes/cells from the same individual). In this section, we analyzed the expected occurrence of redundant CO identification as a result of a limited number of haploid genomes in a pool. For this we refer to the number of recombinant molecules that result from the same CO event as *molecule coverage* of a CO event. To model the expected distribution of *molecule coverages* in a pool we assumed that the recombinant molecules would be distributed among CO events following a Poisson distribution.

Based on the total number of distinct CO events *N* and the recombinant molecules *n*, the average molecule coverage across all distinct CO events would be *λ*=*n*/*N*. Thus, the expected number of COs covered by *k* molecules (*k*=0, 1, 2, …, *n*) would be *N* ∗ *e*^-^*^λ^* ∗ *λ^k^* / *k*! according to the Poisson distribution.

To estimate the effects of varying numbers of distinct, haploid genomes *G*, we first assumed a total number of distinct CO events *N*, and a recombinant molecule number *n*. *N* was set as *N*=*G* ∗ 4.15 (i.e. the average number of COs per haploid genome). For the recombinant molecule number, we estimated the probability of a molecule to cover a CO as 10 kb ∗ 4.15 / 120Mb (=3.33e-04), assuming a core molecule length (region in which a CO can be identified) of 10 kb, 4.15 CO events per haploid genome and a haploid genome size of 120 Mb. Further assuming a 10X library with 10 million molecules, we finally estimated a recombinant molecule number *n* of 3,333 per library.

Using these two parameters and an increasing number of haploid genomes *G*, we investigated the effect of *G* on the molecule coverages in a pool (Supplementary Table 6). Results showed that with 400 haploid genomes (200 F_2_s), nearly two thirds of the COs would be covered by more than one molecule. In a pool with 2,500 haploid genomes (1,250 F_2_s), already 85% of the recombinant molecules would result from distinct CO events. To further increase this fraction by another 10% to 95%, the number of haploid genomes would need to be increased by more than ~3 fold to 8,000.

*Overlapping recombinant molecules from overlapping CO events*

In addition to recombinant molecules sampled from the same CO event, recombinant molecules can also overlap if the underlying CO events overlap. To estimate the fraction of overlapping recombinant molecules from distinct (but overlapping) CO events, we segmented the reference genome (120Mb) into 4,000 genomic bins of 30 kb.

Assuming that the distribution of COs within the bins follows a Poisson distribution, we calculated the Poisson parameter *λ* by dividing the number of observed COs by the number of bins. For example, there were 2,538/4,000 COs per bin on average for P200R1. The expected number of bins with *k* (0, 1, 2, …) COs was therefore 4,000 ∗ 2.71828^(-*λ*) ∗ *λ*^*k* / *k*!.

The expected number of unique COs in all bins with *k* COs was 2-2/2^*k*, as the CO molecules with the different CO transition patterns (Col-0=>L*er* *vs.* Col-0=>L*er*) could be distinguished even if they overlap. Summarizing the expected number of unique COs for each bin, we estimated the total number of unique COs given an observed CO molecule number (Supplementary Table 5). For instance, for 2,538 predicted CO molecules (in P200R1), there were 14.3% CO molecules that overlapped due to the overlap of the underlying, independent CO molecules. When pool size increases, the random overlap become more frequent. For example, for P1250R2 with 3,430 predicted CO molecules, there were 18.7% CO molecules that overlapped due to the overlap of the underlying, independent CO molecules.

Taken together, within pools with a low number of genomes, CO molecules overlap due to the repeated sampling of identical molecules, whereas in large pools even overlapping CO molecules result from independent CO events (as these overlap). Therefore, it is justified to combine overlapping CO molecules in small pools but treat overlapping CO molecules in large pools as independent CO events.

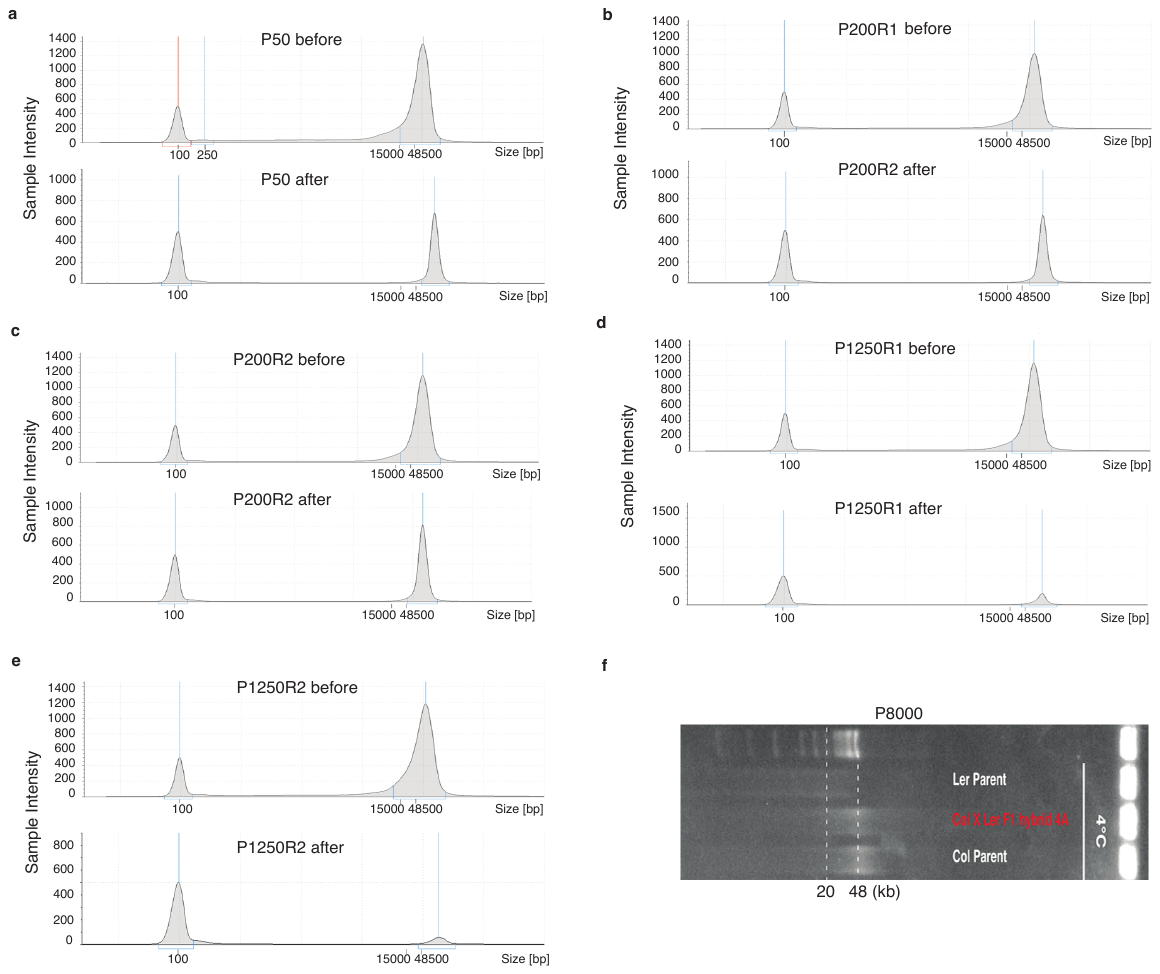

**Supplementary Figure 1. Length of DNA molecules loaded into the Chromium Controller.**

***a-e***. Molecule size distributions of the F_2_ DNA pools before (upper) and after (lower) size selection (by Blue Pippin to obtain molecules longer than 40 kb). The peak at 100 bp is a size marker. ***f***. The F_1_ pollen DNA molecules are distributed around 48 kb.

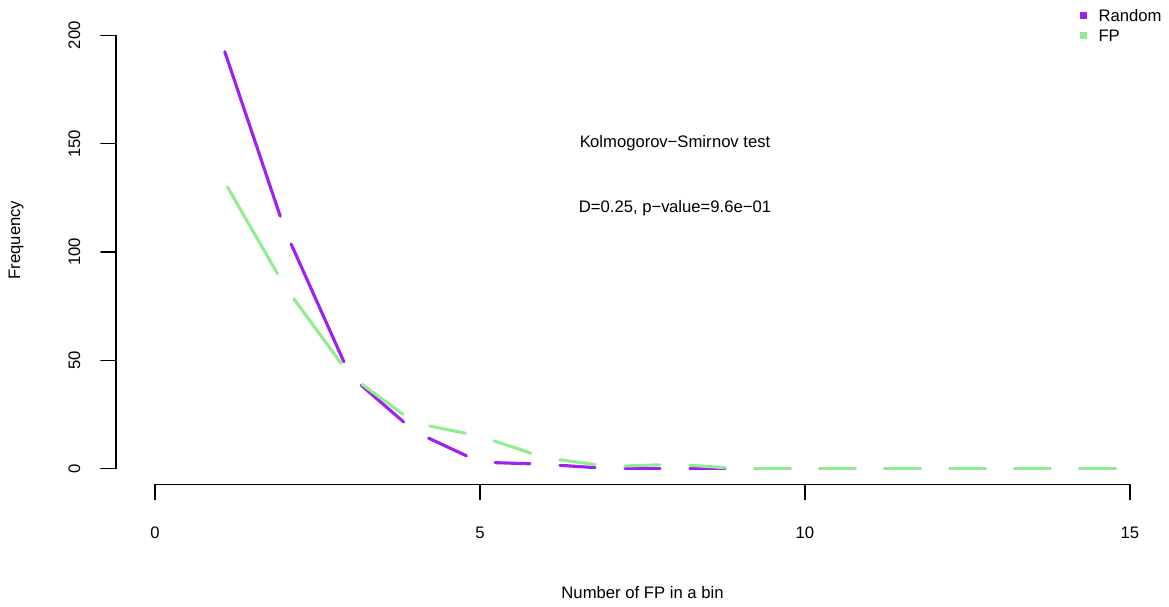

**Supplementary Figure 2. Distribution of false positive CO predictions (P50L75) in genomic bins compared with randomly placed COs.**

The distributions of the 645 false positive CO predictions and 645 randomly placed COs in 4,000 non-overlapping 30 kb bins across the Col-0 reference genome are not significantly different from each other according to a *Kolmogrov-Smirnov* test (*p*-value=9.6e-01, *ks.boot* function in *R* package “Matching”), suggesting that false positives are mostly randomly distributed.

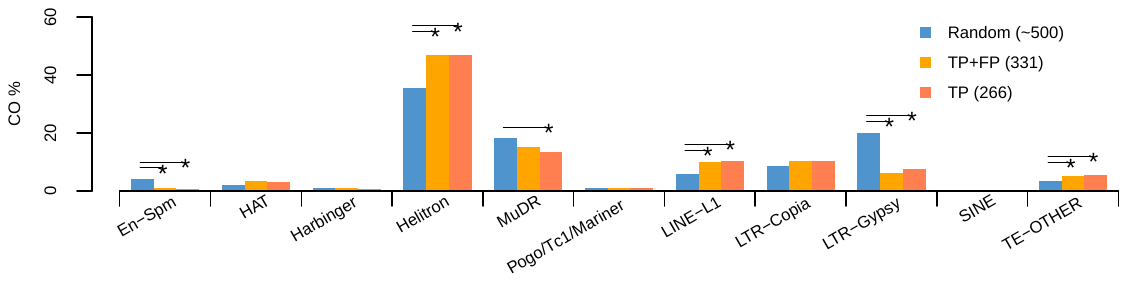

**Supplementary Figure 3. CO frequency within transposable elements (TE) super families.**

A random overlap expectation was estimated by randomly distributing CO-sized intervals within the reference genome (excluding heterochromatic regions). The middle position of each interval was used to assess the overlap with a TE. For each TE family, a permutation test (oneway_test in *R* “coin” package) was performed between the TP or TP+FP sets with 1,000 iterations of randomly inserted CO intervals (each with ~5,000 random intervals). The asterisks indicate significant differences between random sampling and observed overlaps. Most of the significant differences that were present between the random set and the true CO sites, could also be found with the entire set of COs (TP+FP). TP: true positive; FP: false positive.

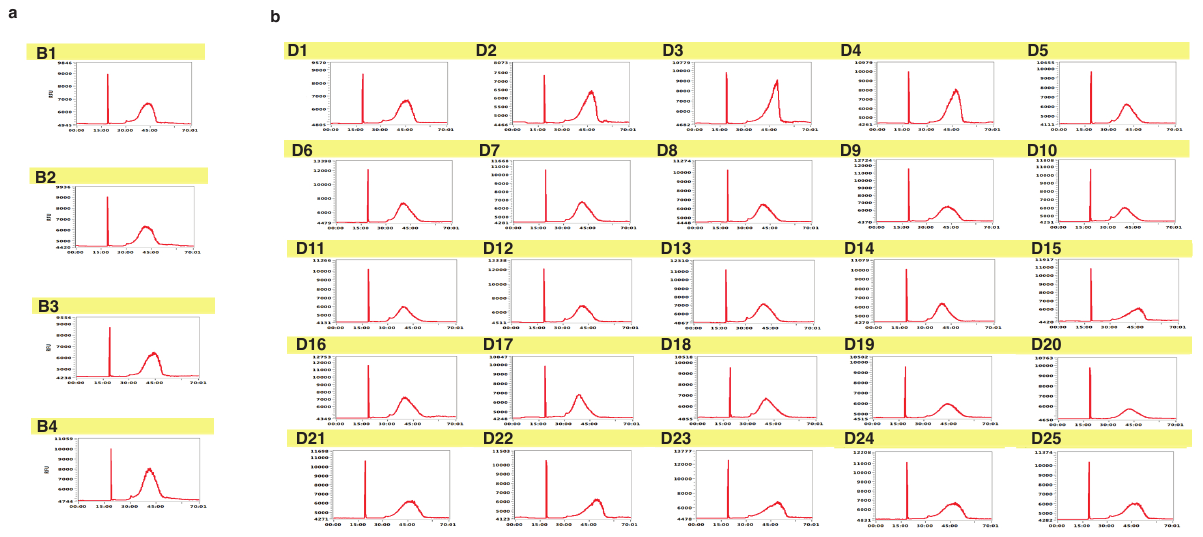

**Supplementary Figure 4. Molecule size distribution (FEMTOpulse, AATI) for each of the 25 50-F_2_ pools.**

***a*.** DNA pools B1-B4 (each generated from DNA extractions of bulked leaves material of 50 F_2_ plants) showed comparable molecule size distributions and were pooled with equal concentration to create libraries P200R1 and P200R2. ***b***. DNA pools D1-D25 (again each from 50 F_2_s) showed some variation in their molecule size distributions and were therefore pooled with equal molarity in respect to the molecules, which were 42~70 kb in size to create libraries P1250R1 and P1250R2. Note: *x*-axis: size of molecules (kb), *y*-axis: RFU - Relative fluorescence unit.

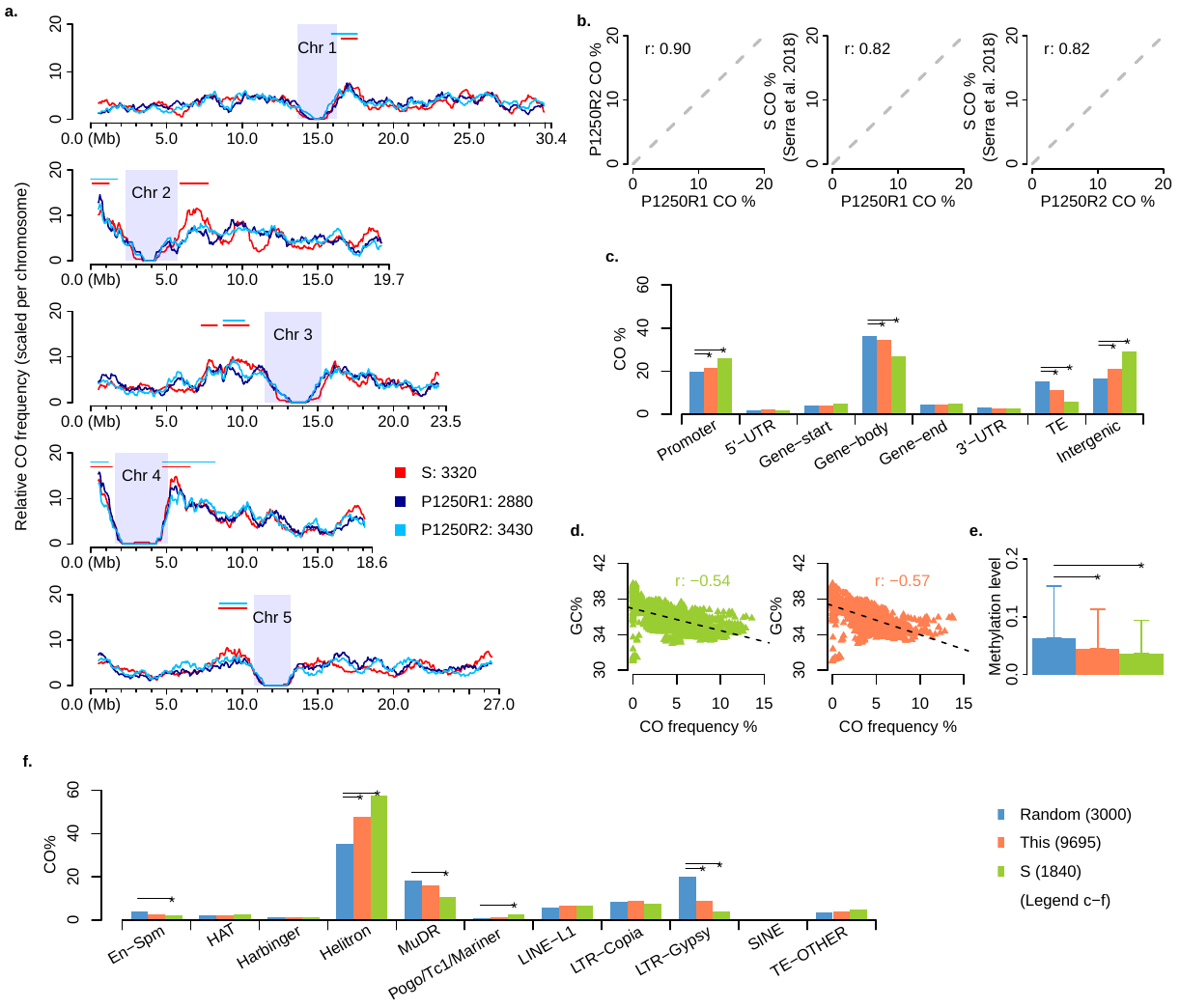

### Supplementary Figure 5. High consistency of predicted CO patterns with CO sites assessed in a different population generated from the same parental lines^4^. *a.* Sliding window-based genome-wide CO landscapes of P1250R1/2 and the dataset S, which refers to 3,320 CO sites detected using whole-genome sequencing of 437 individual plants selected from different F_2_ populations generated from the same parental lines^4,5^, where *frequency* for a window is CO count in the window divided by total CO count of the respective chromosome. Heterochromatic regions^6^ are indicated by rectangles in light blue. Horizontal lines indicate hot regions in CO occurrence that were nearly consistently identified using P1250R2 and S (with comparable numbers of COs), where a hot region was merged from the 2.5% windows with the highest CO frequency while *frequency* for a window was CO count in the window divided by the total CO count of the library. *b.* Correlation of CO frequency (corresponding to windows in *a*). *c.* Association of 9,695 COs (a union set of COs of P200R1/R2 and P1250R1/R2, S^4^) with genomic features, with comparison to a random expectation. Note that compared with Fig. 3c (main text), the inconsistent test at gene end does not exist anymore with the help of a sufficient amount of COs. *d.* Correlations of GC-content with CO frequency (corresponding to windows in *a*). *e.* Methylation in CO intervals, with comparison to a random expectation. *f*. Frequency of COs in transposable element (TE) super families, with comparison to a random expectation. Although the 9,695 COs did not agree with S on the significant overlap with En-Spm, MuDR and Pogo/Tc1/Mariner, these elements made up only ~2% of the total COs. Note: all values were calculated in the same way as given by Fig. 3, Materials and Methods (Main text) and Supplementary Figure 3.

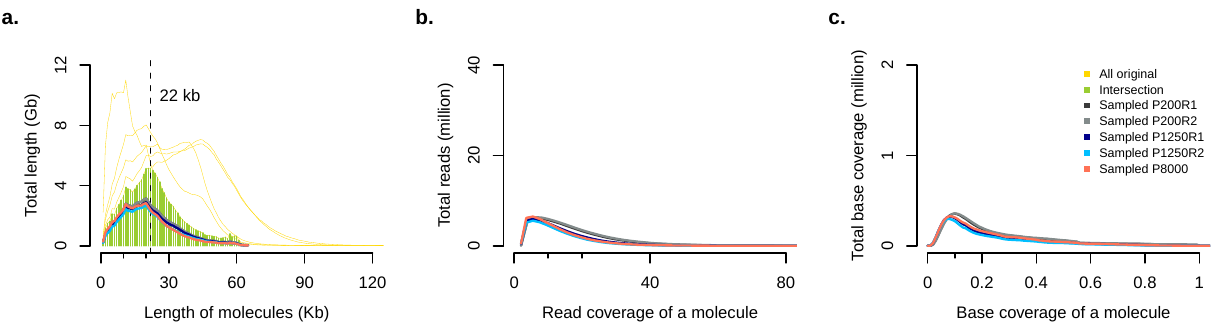

**Supplementary Figure 6. Illustration of molecule characteristics after subsampling.**

***a.*** Length of molecules after subsampling. Lines in gold are the original distributions of P200R1/R2, P1250R1/R2 and P8000, of which the intersection is shown by the solid background (light green) defining the target distribution of the subsampling (Main text: Materials and Methods). ***b.*** Number of reads per molecule after subsampling. ***c.*** Molecule base coverage after subsampling.

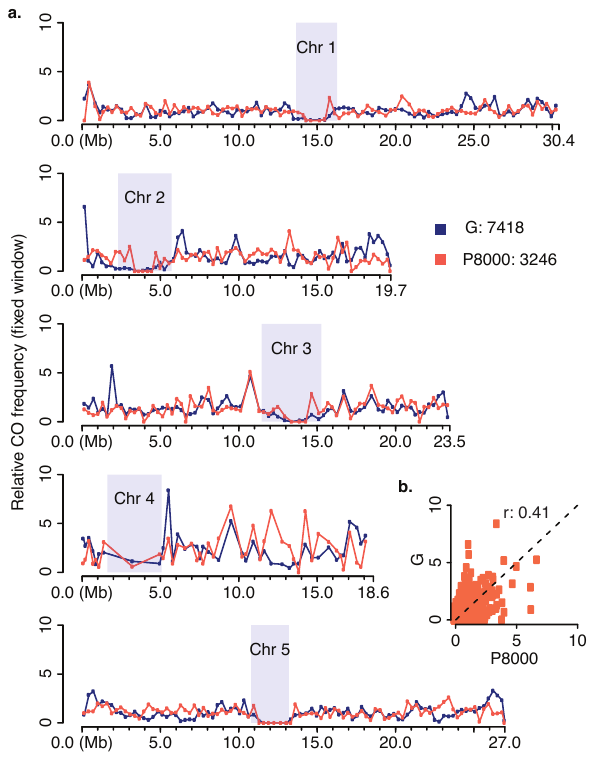

### Supplementary Figure 7. Landscape of COs detected in pollen is consistent to COs introduced by male meiosis and identified in a Col-0 x L*er* backcross population^6^. CO frequency along chromosomes were calculated within reported windows^6^ at an average marker distance of 316.4 kb. Heterochromatic regions^6^ are indicated by rectangles in gray. Although local differences were observed especially at the end of chromosomes, there was a significant genome-wide consistency between the two analyses (Pearson’s *r* 0.41, *p*-value < 2.2e-16). Note: to minimize the effect of these large fixed windows, for each window, the CO frequency detected in pollen were corrected by multiplying the ratio of the marker density in the window to the marker density along a chromosome, where the marker density was defined as the length of a region (or chromosome) divided by the number of markers it harbored.

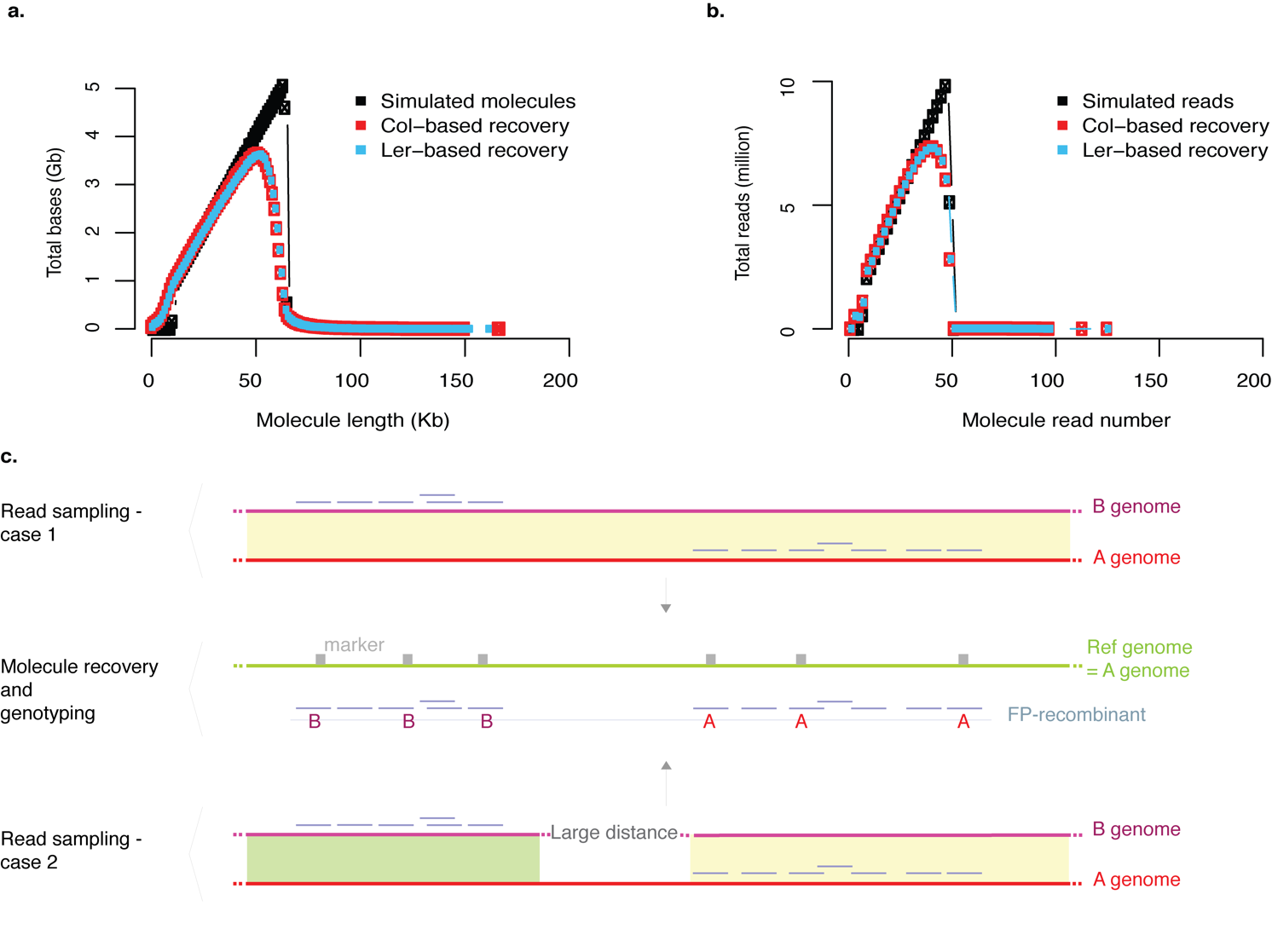

### Supplementary Figure 8. Molecule recovery in simulation and the effect of structural variation.

### *a*. Length distributions of simulated and recovered molecules. *b.* Read distributions of simulated and recovered molecules. Molecule recovery with both Col-0^1^ (red) and L*er*^2^ (light blue) reference genomes are compared with the simulated settings (black). *c*. Random collisions of molecules and false positive (FP). According to *a* and *b*, recovered molecules longer than 65 kb or with more than 50 reads were chimeras (as such molecules were not simulated). In case 1, the molecules are sampled from genomes *A* and *B* with the same barcode and locate in neighboring regions (indicated by solid light yellow), consequently, the recovered molecule form a chimeric FP. In case 2, molecules are sampled from the distantly located rearranged regions between genomes *A* and *B* (indicated by solid light green), which also results in FP. In particular case 2 gave rise to false positives that are with high molecule coverage.
